# Heat stress prevented the biomass and yield stimulation caused by elevated CO_2_ in two well-watered wheat cultivars

**DOI:** 10.1101/2021.11.21.469459

**Authors:** Sachin G. Chavan, Remko A. Duursma, Michael Tausz, Oula Ghannoum

## Abstract

To investigate the interactive effects of elevated CO_2_ and heat stress (HS), we grew two contrasting wheat cultivars, early-maturing Scout and high-tillering Yitpi, under non-limiting water and nutrients at ambient (aCO_2_, 450 ppm) or elevated (eCO_2_, 650 ppm) CO_2_ and 22°C in the glasshouse. Plants were exposed to two 3-day HS cycles at the vegetative (38.1°C) and/or flowering (33.5°C) stage.

At aCO_2_, both wheat cultivars showed similar responses of photosynthesis and mesophyll conductance to temperature and produced similar grain yield. Relative to aCO_2_, eCO_2_ enhanced photosynthesis rate and reduced stomatal conductance and maximal carboxylation rate (*V_cmax_*). During HS, high temperature stimulated photosynthesis at eCO_2_ in both cultivars, while eCO_2_ stimulated photosynthesis in Scout. Electron transport rate (*J_max_*) was unaffected by any treatment. eCO_2_ equally enhanced biomass and grain yield of both cultivars in control, but not HS, plants. HS reduced biomass and yield of Scout at eCO_2_. Yitpi, the cultivar with higher grain nitrogen, underwent a trade-off between grain yield and nitrogen. In conclusion, eCO_2_ improved photosynthesis of control and HS wheat, and improved biomass and grain yield of control plants only. Under well-watered conditions, HS was not detrimental to photosynthesis or growth but precluded a yield response to eCO_2_.

**Key message:** High temperatures increased photosynthetic rates only at eCO_2_ and photosynthesis was upregulated after recovery from heat stress at eCO_2_ in Scout suggesting that eCO_2_ increased optimum temperature of photosynthesis.

## Introduction

Ongoing climate change is threatening the production of agricultural crops including wheat (*Triticum aestivum*) (Asseng *et al*., 2015; Mishra *et al*., 2021). By the end of this century, atmospheric carbon dioxide concentration ([CO_2_]) is expected to reach 700 ppm, increasing surface temperatures by 1.1°C to 2.6°C (IPCC, 2014). For every degree of the temperature increase, global wheat production is predicted to decrease by 6–10% (Asseng *et al*., 2015; García *et al*., 2015). Crop models are important tools for assessing the impact of climate change (Asseng *et al*., 2013). However, they largely lack the ability to consider genotype-specific responses to elevated [CO_2_] (eCO_2_) and their interaction with other environmental conditions. Hence, it is important to better understand how plants respond to eCO_2_ interactions with the environment. Photosynthesis, a fundamental process driving crop growth and yield, can partially explain the interactive effects of eCO_2_ with environmental stresses and provide a mechanistic basis for crop models (Yin and Struik, 2009).

During photosynthesis, ribulose-1, 5-bisphosphate carboxylase (Rubisco) catalyzes the carboxylation and oxygenation of ribulose-1, 5-bisphosphate (RuBP). eCO_2_ increases photosynthetic rates (*A_sat_*) and reduces photorespiration and stomatal conductance (*g_s_*). Generally, higher photosynthetic rates enhance the growth and productivity of plants leading to increased leaf area, plant size and crop yield (Krenzer and Moss, 1975; Sionit *et al*., 1981; Hocking and Meyer, 1991; Mitchell *et al*., 1993; Kimball *et al*., 1995; Mulholland *et al*., 1998; Cardoso–Vilhena and Barnes, 2001; Högy *et al*., 2009; Kimball, 2016; Fitzgerald *et al*., 2016; Kimball, 1983). Following long term CO_2_ enrichment, photosynthetic capacity may diminish due to lower amount of Rubisco (Nie *et al*., 1995; Rogers and Humphries, 2000; Ainsworth *et al*., 2003) or reduced activation of Rubisco (Delgado *et al*., 1994).

Optimum temperature range for wheat growth is 17-23°C, with a minimum of 0°C and maximum of 37°C (Porter and Gawith, 1999). Global warming involves a gradual increase in mean temperature as well as increased frequency and intensity of heat waves. Heat can adversely affect crop growth and disrupt reproduction depending on the timing, intensity and duration (Sadras and Dreccer, 2015). Higher daytime temperatures (below damaging level) increase photosynthesis up to an optimum temperature, above which photosynthesis decreases mainly due to higher photorespiration (Berry and Bjorkman, 1980; Long, 1991). High night time temperatures increase respiration and reduce overall photosynthetic carbon gain (Prasad *et al*., 2008). At the whole plant level, high temperatures accelerate growth (Fischer, 1980) and shorten crop duration (Hatfield and Prueger, 2015), hence reducing grain yield due to insufficient time to capture resources. Losses due to short crop duration are usually higher than benefits of growth stimulation at high temperature (Wardlaw and Moncur, 1995).

The severity of the damage caused by abrupt temperature increases above the optimum range (termed heat stress, HS) depends on the magnitude and duration of HS as well as the developmental stage of the plant (Wahid et al., 2007). HS may reduce photosynthesis due to reduced chlorophyll content, impaired photosystem II and lower Rubisco activation (Berry and Bjorkman, 1980; Eckardt and Portis, 1997). HS can directly damage cells and increase grain abortion resulting in reduced growth, biomass and grain yield (Stone and Nicolas, 1996, 1998; Wardlaw *et al*., 2002; Farooq *et al*., 2011). Around anthesis, HS (>30°C) reduces seed setting due to lower pollen viability, leading to poor fertilization, abnormal ovary development and slower pollen growth (Balla *et al*., 2019).

The interactive effects of eCO_2_ and HS can be positive, negative or neutral (Wang et al., 2008, 2011). Elevated CO_2_ increases the temperature optimum of photosynthesis (Long, 1991; Alonso *et al*., 2009) by reducing photorespiration and improving tolerance to photoinhibition (Hogan *et al*., 1991). The impact of HS on photosynthesis will depend on whether Rubisco, electron transport or end-product synthesis is limiting at eCO_2_ (Sage and Kubien, 2007). Enhanced growth and leaf level water use efficiency (WUE) by eCO_2_ may help compensate for the negative impact of HS; conversely, heat-induced shortening of the grain-filling stage and grain abortion could limit the benefits of eCO_2_ (Lobell and Gourdji, 2012). In addition, decreased *g_s_* under eCO_2_ may limit transpirational cooling and therefore exacerbate HS. Thus, HS counteracts the positive effect of eCO_2_ on yield components and may aggravate the negative effect of eCO_2_ on grain quality due to the high sensitivity of wheat to temperature stress especially during anthesis and grain-filling stage (Wang & Liu, 2021).

Many studies have investigated the response of wheat to eCO_2_ in enclosures and in the field (Wang & Liu, 2021). However, only a few studies have considered eCO_2_ interaction with temperature increases in wheat (Rawson, 1992; Delgado *et al*., 1994; Morison and Lawlor, 1999; Jauregui *et al*., 2015; Cai *et al*., 2016) and rarely with HS (Coleman *et al*., 1991; Wang *et al*., 2008). Studies considering HS have addressed mainly the biomass or yield aspects and not the physiological processes such as photosynthesis (Stone and Nicolas, 1994, 1996, 1998). Interactive effects of eCO_2_ and HS on photosynthesis have been reported in a limited number of studies (Wang *et al*., 2008, 2011; Macabuhay, 2016; Macabuhay *et al*., 2018; Chavan *et al*., 2019). Macabuhay et al., (2018) studied interactive effects of eCO_2_ and (experimentally imposed) heatwaves on wheat (cv Scout and Yitpi) grown in a dryland cropping system and concluded that eCO_2_ may moderate some effects of HS on grain yield but such effects strongly depend on seasonal conditions and timing of HS. In another glasshouse experiment on the interactive effects of severe HS and eCO_2_ in wheat (cv Scout), we found that eCO_2_ mitigated the negative impacts of HS at anthesis on photosynthesis and biomass, but grain yield was reduced by HS in both CO_2_ treatments (Chavan *et al*., 2019). However, HS can occur throughout plant growth, including during vegetative, flowering or grain filling stages. In addition, different crop genotypes may respond variably to the interaction of eCO_2_ with HS.

Here, we build on our previous work by comparing the interactive effects of eCO_2_ and HS in two commercial wheat cultivars. Scout and Yitpi have similar genetic background but distinct agronomic features. Scout is a mid-season maturity cultivar with very good early vigor that can produce leaf area early in the season. Scout has a putative water-use efficiency (WUE) gene, which has been identified using carbon isotope discrimination (Condon *et al*., 2004). Yitpi is a good early vigor, freely tillering, late flowering and long maturity cultivar (Bahrami et al., 2017; Pacificseeds, 2009; Seednet, 2005).

Although Scout is known to be a high yielding variety with very good grain quality and high reproductive sink (Pacific seeds, 2009), we hypothesized that Yitpi will produce higher grain yield due to its ability to initiate more tillers and its longer time to flower and mature (Hypothesis 1). Fast growing plants with high sink capacity show a greater eCO_2_-induced growth stimulation (Poorter, 1993) and less photosynthetic acclimation (Delgado *et al*., 1994) compared to slow growing counterparts with low sink capacity. Consequently, we hypothesized that Yitpi will show greater photosynthetic, growth and yield response to eCO_2_ due to its greater vegetative sink capacity (tillering) relative to Scout with restricted tillering (Hypothesis 2). The greater growth stimulation at eCO_2_ may buffer Yitpi against HS damage compared to Scout. Thus, HS may decrease yield in Scout more than Yitpi and aCO_2_ more than eCO_2_ (Hypothesis 3). HS is more damaging at the reproductive relative to the vegetative developmental stage (Farooq *et al*., 2011). Hence, we expect less damage in plants exposed to HS at the vegetative stage relative to the flowering stage (Hypothesis 4).

To test these hypotheses, Scout and Yitpi were grown at ambient or elevated CO_2_ conditions and subjected to one or two heat stresses at the vegetative (HS1) and/or flowering (HS2) stage. Growth, biomass and photosynthetic parameters were measured at different time points across the life cycle of the plants. At the plant level, we report that eCO_2_ equally stimulated grain yield of Scout and Yitpi. While moderate HS under well-watered conditions was not detrimental to photosynthesis and growth in the long term due to the transpirational cooling, HS prevented the wheat plant from reaping the benefits of eCO_2_ on biomass and yield.

## Materials and methods

### Plant culture and treatments

The experiment was conducted in the glasshouse facility located at the Hawkesbury campus of Western Sydney University (WSU). Seeds of commercial winter wheat cultivars Scout and Yitpi were procured from Agriculture Victoria (Horsham). Cultivars were selected based on their use in the Australian Grains Free Air CO_2_ Enrichment (AGFACE) project investigating climate change impacts on wheat growth and yield (Houshmandfar *et al*., 2017). For germination, 300 seeds of each cultivar were sterilized using 1.5 % NaOCl_2_ for 1 min followed by incubation in the dark at 28°C for 48 hours in petri plates. Sprouted seeds were planted in germination trays using seed raising and cutting mix (Scotts, Osmocote^®^) at ambient CO_2_ (aCO_2_, 400 μl L^-1^), temperature (22/14 °C day/night), relative humidity (RH, 50 to 70%) and natural light (midday average 500 μmol m^-2^ s^-1^) (Figure S1). The growth stages are denoted by decimal code (DC) according to (Zadoks *et al*., 1974) along with the time points here after. Two-week-old seedlings (DC12) were transplanted to individual cylindrical pots (15 cm diameter and 35 cm height) using sieved soil collected from local site. At transplanting stage (T0) pots were distributed into two aCO_2_ (400μl L^-1^ and two eCO_2_ (650 μl L^-1^) chambers (Figure S1B). Some plants were exposed to one or two HS cycles at the vegetative (HS1, 10 weeks after planting, WAP, DC 32) and/or the flowering (HS2, 15 WAP, DC 63) stages for 3 days with temperature ramp up from 14°C night temperature (8 pm to 6 am) to 38°C during mid-day (10 pm to 4 pm) at 60% daytime RH (Figures S1 and S2). The two HS cycles created four sets of heat treatments at each CO_2_ concentration as follows: (1) Control – plants were not exposed to HS at any stage, (2) HS1 – plants were exposed to HS at vegetative (DC32) stage only, (3) HS2 – plants were exposed to HS at reproductive (DC63) stage only and (4) HS1+2 – plants were exposed to both the heat stresses HS1 and HS2 (Figure S2).

Thrive all-purpose fertilizer (Yates) was applied monthly throughout the experiment to maintain similar nutrient supply in all treatment combinations. Pots were regularly swapped between left and right benches as well as between front and back for randomization within chamber. Pots and treatments were also swapped between the two ambient and two elevated CO_2_ chambers for randomization among chambers.

### Growth and biomass measurements

The full factorial experimental design included four chambers (two chambers for each CO_2_ treatment) and five destructive harvests at time points T0 (2 WAP, DC12), T1 (6 WAP, DC28), T2 (10 WAP, DC35), T3 (17, DC65) and T4 (25 WAP, DC90) (Figure S2). Ten plants per treatment per cultivar were measured and harvested at each time point. Morphological parameters were measured followed by determinations of root, shoot and leaf dry mass. Samples were dried for 48 hours in the oven at 60°C immediately after harvesting. Leaf area was measured at time point T1, T2 and T3 using a leaf area meter (LI-3100A, LI-COR, Lincoln, NE, USA). Plant height, leaf number, tiller number and ear (grain inflorescence) number along with developmental stage information (booting (DC45), half-emerged (DC55) or fully emerged (DC60)) were recorded at time points T2 and T3).

### Leaf gas exchange measurements

The youngest fully developed leaf (which was the flag leaf at T3) was used to measure gas exchange parameters. Instantaneous steady state leaf gas exchange measurements were performed at time points T1, T2 and T3 using a portable open gas exchange system (LI-6400XT, LI-COR, Lincoln, USA) to measure light-saturated (photosynthetic photon flux density (PPFD) =1500 µmol m^-2^ s^-1^) photosynthetic rate (*A_sat_*), stomatal conductance (*g_s_*), ratio of intercellular to ambient CO_2_ (*C_i_*/*C_a_*), leaf transpiration rate (E), dark respiration (*R_d_*) and dark- and light-adapted chlorophyll fluorescence (Fv/Fm and Fv′/Fm′, respectively). Dark adapted leaf measurements were conducted by switching off light for 15 minutes. Steady state leaf gas exchange measurements were also performed during and after heat shock along with recovery stage. Plants were moved to a neighboring chamber with ambient CO_2_ levels for short time (20-30 min for each plant) where air temperature was separately manipulated to achieve the desired leaf temperature. The LI-COR 6400-40 leaf chamber fluorometer (LCF) was used to measure gas exchange at a PPFD of 1500 μmol m^-2^ s^-1^ at two CO_2_ concentrations (400 and 650 μl L^-1^) and two leaf temperatures (25 and 35 °C). Photosynthetic down regulation or acclimation was examined by comparing the measurements at common CO_2_ (ambient and elevated CO_2_ grown plants measured at 400 μl L^-1^ CO_2_ partial pressure) and growth CO_2_ (aCO_2_ grown plants measured at 400μl L^-1^ CO_2_ partial pressure and eCO_2_ grown plants measured at 650 650 μl L^-1^ CO_2_ partial pressure).

Dark respiration (*R_d_*) was measured after a dark adaptation period of 15 minutes. Photosynthetic water use efficiency (PWUE) was calculated as *A_sat_* (μmol m^-2^ s^-1^)/ *g_s_* (mol m^-2^ s^-1^). The response of *A_sat_* to variations in sub-stomatal CO_2_ mole fraction (*Ci*) (A-Ci response curve) was measured at T3 in 8 steps of CO_2_ concentrations (50, 100, 230, 330, 420, 650, 1200 and 1800 μl L^-1^) at leaf temperature of 25°C. Measurements were taken around mid-day (from 10 am to 3 pm) on attached last fully expanded or flag leaves of the main stems. Before each measurement, the leaf was allowed to stabilize for 10-20 minutes until it reached a steady state of CO_2_ uptake and stomatal conductance. Ten replicate plants per treatment were measured.

### Mesophyll conductance and temperature response

Mesophyll conductance (*g_m_*) was determined by concurrent gas exchange and stable carbon isotope measurements using portable gas exchange system (LI-6400-XT, LI-COR, Lincoln, NE, USA) connected to a tunable diode laser (TDL) (TGA100, Campbell Scientific, Utah, USA) for the two wheat cultivars grown at ambient atmospheric CO_2_ partial pressures. *A_sat_* and ^13^CO_2_/^12^CO_2_ carbon isotope discrimination were measured after T1 at five leaf temperatures (15, 20, 25, 30 and 35°C) and saturating light (1500 µmol quanta m^-2^ s^-1^). Leaf temperature sequence started at 25°C decreasing to 15°C and then increased up to 35°C. Response of *A_sat_* to variations in *Ci* was measured at each leaf temperature. Dark respiration was measured by switching light off for 20 minutes at the end of each temperature curve. Measurements were made inside a growth cabinet (Sanyo) to achieve desired leaf temperature. The photosynthetic carbon isotope discrimination (Δ) to determine *g_m_* was measured as follows (Evans *et al*., 1986):

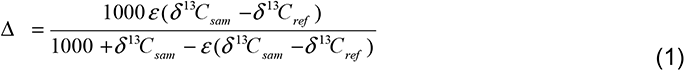

Where,

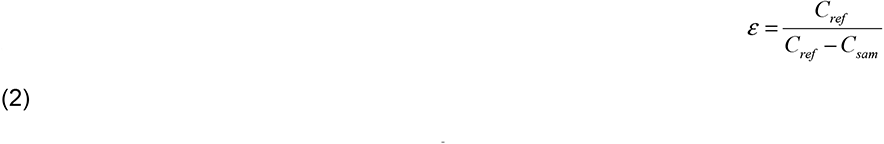

*C*_ref_ and *C*_sam_ are the CO_2_ concentrations of dry air entering and exiting the leaf chamber, respectively, measured by the TDl. *g_m_* was calculated using correction for ternary and second-order effects (Farquhar and Cernusak, 2012; Evans and Von Caemmerer, 2013) following the next expression:

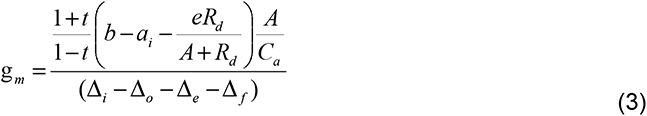

Where, Δ_i_ is the fractionation that would occur if the *g*m were infinite in the absence of any respiratory fractionation (*e* = 0), Δ_o_ is observed fractionation, Δ_e_ and Δ_f_ are fractionation of ^13^C due to respiration and photorespiration respectively (Evans and Von Caemmerer, 2013).

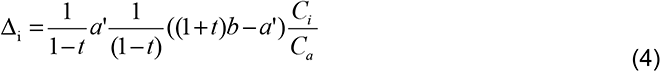

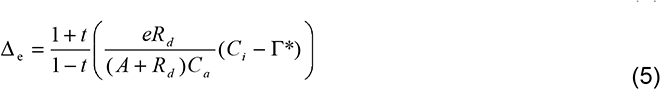

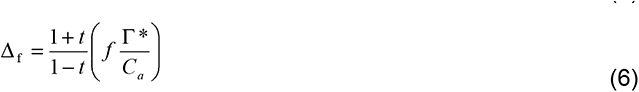

Where,

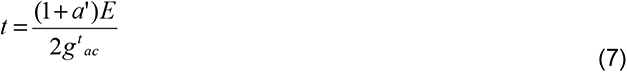

The constants used in the model were as follows: E denotes transpiration rate; g^t^_ac_ is total conductance to diffusion in the boundary layer (*a*b = 2.9‰) and in air (*a* = 4.4‰); *a’* is the combined fractionation of CO_2_ across boundary layer and stomata; net fractionation caused by RuBP and PEP carboxylation (*b* = 27.3‰) (Evans *et al*., 1986); fractionation with respect to the average CO_2_ composition associated with photorespiration (*f* = 11.6‰) (Lanigan *et al*., 2008) and we assumed null fractionation associated with mitochondrial respiration in light (*e* = 0).

### Leaf nitrogen and carbon estimation

Leaf discs were cut from the flag leaves used for gas exchange measurements at time points T2 and T3 then oven dried. Leaf discs were processed for nitrogen (N) and carbon (C) content using elemental analyzer (Dumas method). N and C were also estimated from other plant components including leaf, stem, root and grain harvested at T1, T3 and T4. Ground samples were processed for C & N with a CHN analyzer (LECO TruMac CN-analyser, Leco corporation, USA) using an automated dry combustion method (Dumas method). Leaf N per unit area (N_area_) was calculated as N (mmol g^-1^) × LMA (g m^-2^). Photosynthetic nitrogen use efficiency (PNUE) was calculated as *A_sat_* (μmol m^-2^ s^-1^)/leaf N_area_ (mmol m^-2^). Protein content was determined using N and multiplication factor of 5.7 (Mosse, 1990; Bahrami *et al*., 2017).

### Statistical and temperature analysis

All data analyses and plotting were performed using R computer software (R Core Team, 2020). The effect of treatments and their interactions was analyzed using linear modeling with ‘anova’ in R. Significance tests were performed with anova and post hoc Tukey test using the ‘glht’ function in the multcomp R package. Coefficient means were ranked using post-hoc Tukey test. The Farquhar-von Caemmerer-Berry (FvCB) photosynthesis model was fit to the *A_sat_* response curves to C_i_ (A-Ci response curve) or chloroplastic CO_2_ mole fraction (Cc), which was estimated from the *g_m_* measurements (A-Cc response curve). We used the plantecophys R package (Duursma, 2015) to perform the fits, using measured *g_m_* and *R_d_* values, resulting in estimates of maximal carboxylation rate (*V_cmax_*) and maximal electron transport rate (*J_max_*) for D-ribulose-1,5-bisphosphate carboxylase/oxygenase (Rubisco) using measured *R_d_* values. Temperature correction parameter (Tcorrect) was set to False while fitting A-C_i_ curves. Temperature response of *V_cmax_* and *J_max_* were calculated by Arrhenius and peaked functions, respectively (Medlyn *et al*., 2002). Estimated *V_cmax_* and *J_max_* values at five leaf temperatures were then fit using nonlinear least square (nls) function in R to determine energy of activation for *V_cmax_* (EaV) and *J_max_* (EaJ) and entropy (ΔSJ). Temperature responses of *V_cmax_* and R_d_ were fit using Arrhenius equation as follows,

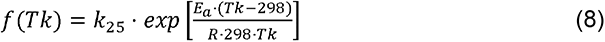

Where, Ea is the activation energy (in J mol^-1^) and k_25_ is the value of R_d_ or *V_cmax_* at 25 °C. R is the universal gas constant (8.314 J mol^-1^ K^-1^) and Tk is the leaf temperature in K. The activation energy term Ea describes the exponential rate of rise of enzyme activity with the increase in temperature. The temperature coefficient Q_10_, a measure of the rate of change of a biological or chemical system as a consequence of increasing the temperature by 10 °C was also determined for *Rd* using the following equation:

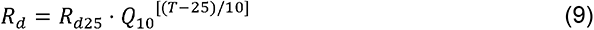

A peaked function (Harley *et al*., 1992) derived Arrhenius function was used to fit the temperature dependence of *J_max_*, and is given by the following equation:

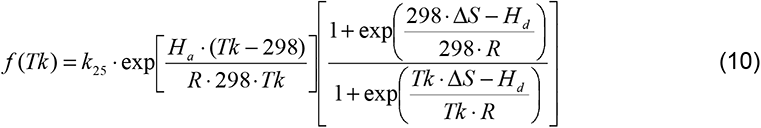

Where, Ea is the activation energy and k_25_ is the *J_max_* value at 25 °C, Hd is the deactivation energy and S is the entropy term. Hd and ΔS together describe the rate of decrease in the function above the optimum. Hd was set to constant 200 kJ mol^-1^ to avoid over parametrization. The temperature optimum of *J_max_* was derived from Eqn 10 (Medlyn *et al*., 2002) and written as follows:

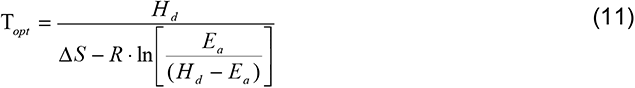

The temperature response of *A_sat_* was fit using a simple parabola equation (Crous *et al*., 2013) to determine temperature optimum of photosynthesis:

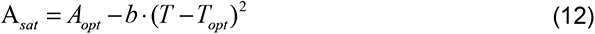

Where, T is the leaf temperature of leaf gas exchange measurement for *A_sat_*, *T_opt_* represents the temperature optimum and *A_opt_* is the corresponding *A_sat_* at that temperature optimum. Steady state gas exchange parameters *g_m_*, *g_s_, C_i_* and *J_max_* to *V_cmax_* ratio were fit using nls function with polynomial equation:

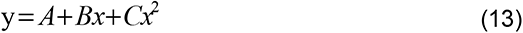

## Results

Two commercial wheat cultivars Scout and Yitpi were grown under aCO_2_ or eCO_2_ (daytime average of 450 or 650 μl L, respectively; 65% RH and 22°C), natural sunlight and well-watered conditions (Figure S1). Both aCO_2_ and eCO_2_ grown plants were exposed to two 3-day HS cycles at the vegetative (HS1, 10 WAP, DC32, daytime average of 38°C) and flowering stage (HS2, 15 WAP, DC63, daytime average of 33.5°C), while daytime RH was maintained at 60%. HS2 was lower in intensity relative to HS1 due to the cool winter conditions. Both HS cycles had similar overall effects on growth and yield parameters, refuting our fourth hypothesis that HS during the reproductive stage is more damaging. Hence, we mostly compare the control plants to those exposed to both heat stresses. Grain filling started 17 WAP (DC65) and final harvest occurred 25 WAP (DC90) (Figure S2).

### Photosynthetic temperature responses of the two wheat cultivars at aCO_2_

A-C_i_ curves together with *g_m_* were measured at five leaf temperatures to characterize the thermal photosynthetic responses of the two wheat cultivars grown at aCO_2_ (Figure 1; Table 1). Overall, both cultivars had similar photosynthetic temperature responses. *A_sat_* and *g_s_* increased with leaf temperature up to an optimum (*T_opt_*) around 23.4°C and decreased thereafter, while C_i_ slowly decreased with temperature. Mesophyll conductance (*g_m_*) increased up to 25°C then plateaued (Figure 1B). The temperature response of *g_m_* as well as the values recorded at 25°C (0.25-0.31 mol m^-2^ s^-1^ bar^-1^) for Scout and Yitpi (Figure 1B) were similar to what has been reported for wheat (0.39) and other crop species (cotton = 0.73, soybean = 0.49 and rice = 0.67) (Von Caemmerer and Evans, 2015; Jahan *et al*., 2021). Scout had slightly higher *A_sat_*, *g_s_* and *g_m_* than Yitpi at most leaf temperatures (Figures 1A- D). *R_d_* linearly increased with temperature, and both cultivars had similar Q_10_ (Figure 1H, Table 1). *V_cmax_* and *J_max_* exponentially increased with leaf temperature, but *J_max_* declined above *T_opt_* (30°C) in both cultivars (Figures 1E-F). There was no significant difference in *V_cmax_*, *J_max_* or their activation energies between the two wheat cultivars (Figure 1E-G, Table 1). The ratio of *J_max_*/*V_cmax_* was similar for the two cultivars and linearly decreased with leaf temperature (Figure 1G).

**Figure 1.**
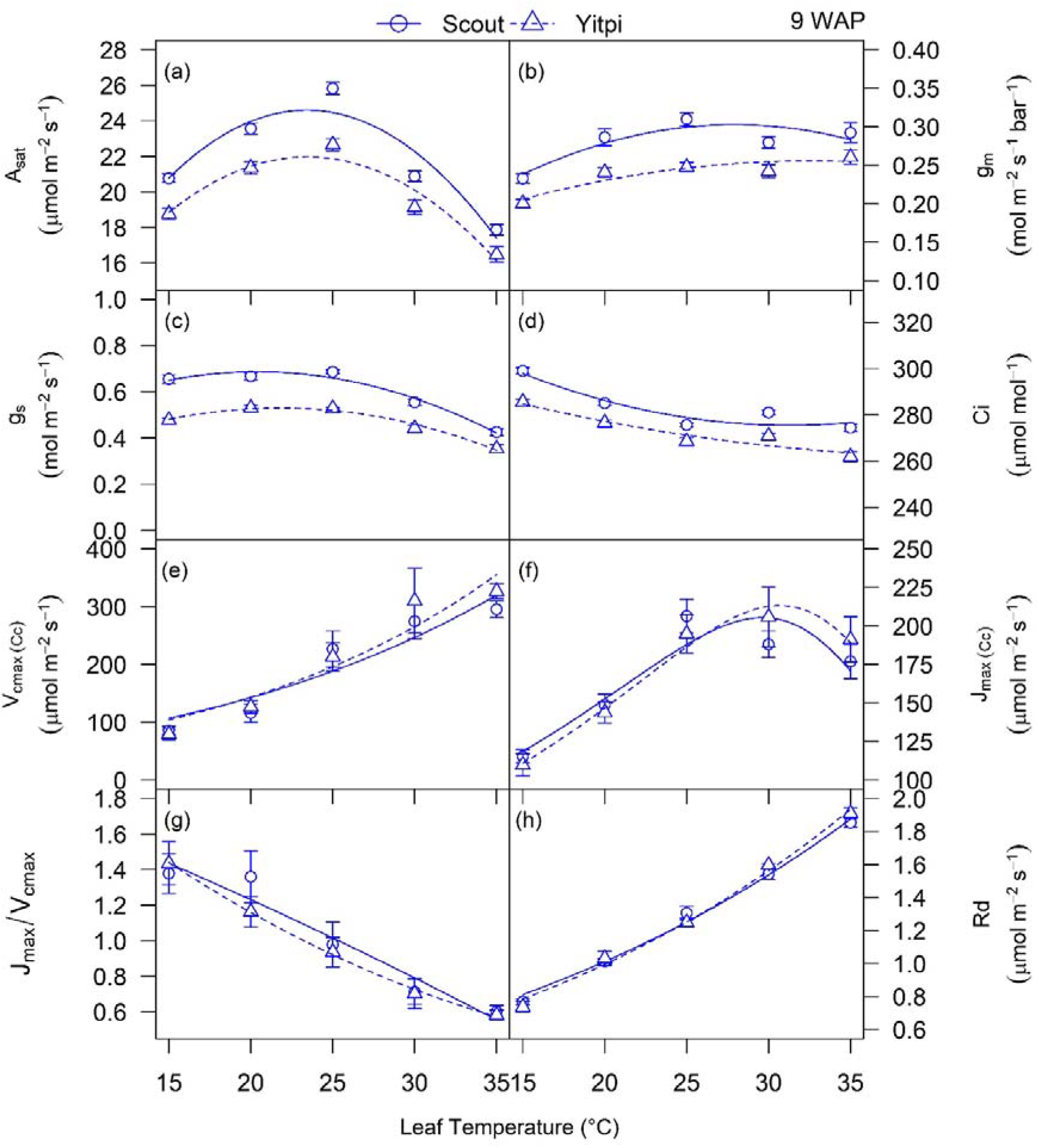
Temperature response of photosynthetic parameters: CO_2_ assimilation rate (a), mesophyll conductance (b), stomatal conductance (c) and intercellular CO_2_ (d), *V_cmax_* (e), *J_max_* (f), *J_max_* / *V_cmax_* (g) and dark respiration (h) over leaf temperatures (15, 20, 25, 30 and 35 °C) in plants grown at aCO_2_. Scout and Yitpi are depicted using circles with solid cultivars and triangles with broken cultivars respectively. Data in panels (a), (b), (c), (d), (e), (f) and (h) are fit using nonlinear least square (nls) function in R.

**Table 1.** Summary of modelled parameters for temperature response of photosynthesis. Summary of coefficients derived using nonlinear least square fitting of CO_2_ assimilation rates and maximal rate of carboxylation (*V_cmax_*) and maximal rate of RuBP regeneration (*J_max_*) determined using A-C_i_ response curves and dark respiration measured at five leaf temperatures 15, 20, 25, 30 and 35°C. Values are means with standard errors. Derived parameters include temperature optima (*T_opt_*) of photosynthesis (A_opt_); activation energy for carboxylation (EaV); activation energy (EaJ), entropy term (ΔSJ) and *T_opt_* and corresponding value for *J_max_* with deactivation energy (Hd) assumed constant; and activation energy (EaR) and temperature coefficient (Q_10_) for dark respiration. Letters indicate significance of variation in means.

| Parameter | Constant | Scout | Yitpi |
| --- | --- | --- | --- |
| **$A_{sat}$**<br>($\mu\text{mol m}^{-2} \text{s}^{-1}$) | $T_{opt}$ (°C) | 23.4 ± 1 a | 23.4 ± 0.7 a |
| | $A_{opt}$ | 24.6 ± 1 a | 22 ± 0.6 b |
| **$g_m$**<br>($\text{mol m}^{-2} \text{s}^{-1} \text{bar}^{-1}$) | $T_{opt}$ (°C) | 27.9 ± 2.2 a | 32.5 ± 6.9 b |
| | $g_m$ at 25°C | 0.31 ± 0.01 a | 0.25 ± 0.01 b |
| | $g_m$ at $T_{opt}$ | 0.30 ± 0.01 | 0.25 ± 0.01 |
| **$V_{cmax}$**<br>($\mu\text{mol m}^{-2} \text{s}^{-1}$) | $V_{cmax}$ at 25°C | 192.7 ± 17.1 a | 198.4 ± 17.7 a |
|  | EaV (kJ mol<sup>-1</sup>) | 43.3 ± 8.74 a | 46.4 ± 8.7 a |
| **$J_{max}$**<br>($\mu\text{mol m}^{-2} \text{s}^{-1}$) | $J_{max}$ at 25°C | 187.9 ± 13.1 a | 186.1 ± 5.7 a |
| | $T_{opt}$ (°C) | 29.6 ± 0.3 a | 30.5 ± 0.3 a |
| | $J_{max}$ at $T_{opt}$ | 205.7 ± 10.2 | 215.4 ± 13.4 |
|  | EaJ (kJ mol<sup>-1</sup>) | 37.7 ± 13.2 a | 41.1 ± 5.8 a |
| | $\Delta SJ$ (J mol<sup>-1</sup> K<sup>-1</sup>) | 648.3 ± 5.3 a | 647 ± 2.4 a |
|  | Hd (kJ mol<sup>-1</sup>) | 200 |  |
| **$Rd$**<br>($\mu\text{mol m}^{-2} \text{s}^{-1}$) | $Rd$ at 25°C | 1.25 ± 0.02 a | 1.25 ± 0.02 a |
|  | EaR (kJ mol<sup>-1</sup>) | 30.9 ± 1.6 a | 33.2 ± 1.7 a |
| | $Q_{10}$ | 1.51 ± 0.03 a | 1.56 ± 0.04 a |

### eCO_2_ stimulated photosynthesis and reduced stomatal conductance in both wheat cultivars

Overall, the two wheat cultivars had similar *A_sat_*, *g_s_*, PWUE (A_sat_/*g_s_*), *R_d_*, Fv/Fm, *V_cmax_* and *J_max_* measured under most conditions (Figures 1-4, Tables S1-S3). Under control (non-HS) conditions, eCO_2_ enhanced *A_sat_* measured at growth CO_2_ (*A_growth_*) and 25℃ and reduced *g_s_* in both cultivars at T1, T2 and T3 (Figure 2, Tables S1-S3). When measured at common CO_2_ and 25°C, eCO_2_-grown plants had lower *A_sat_* (−12% at T2, p <0.01) and *g_s_* (−10% at T2, p <0.001) relative to aCO_2_. This photosynthetic downregulation was more persistent in Yitpi compared to Scout (Figure 2, Tables S1-S3).

**Figure 2.**
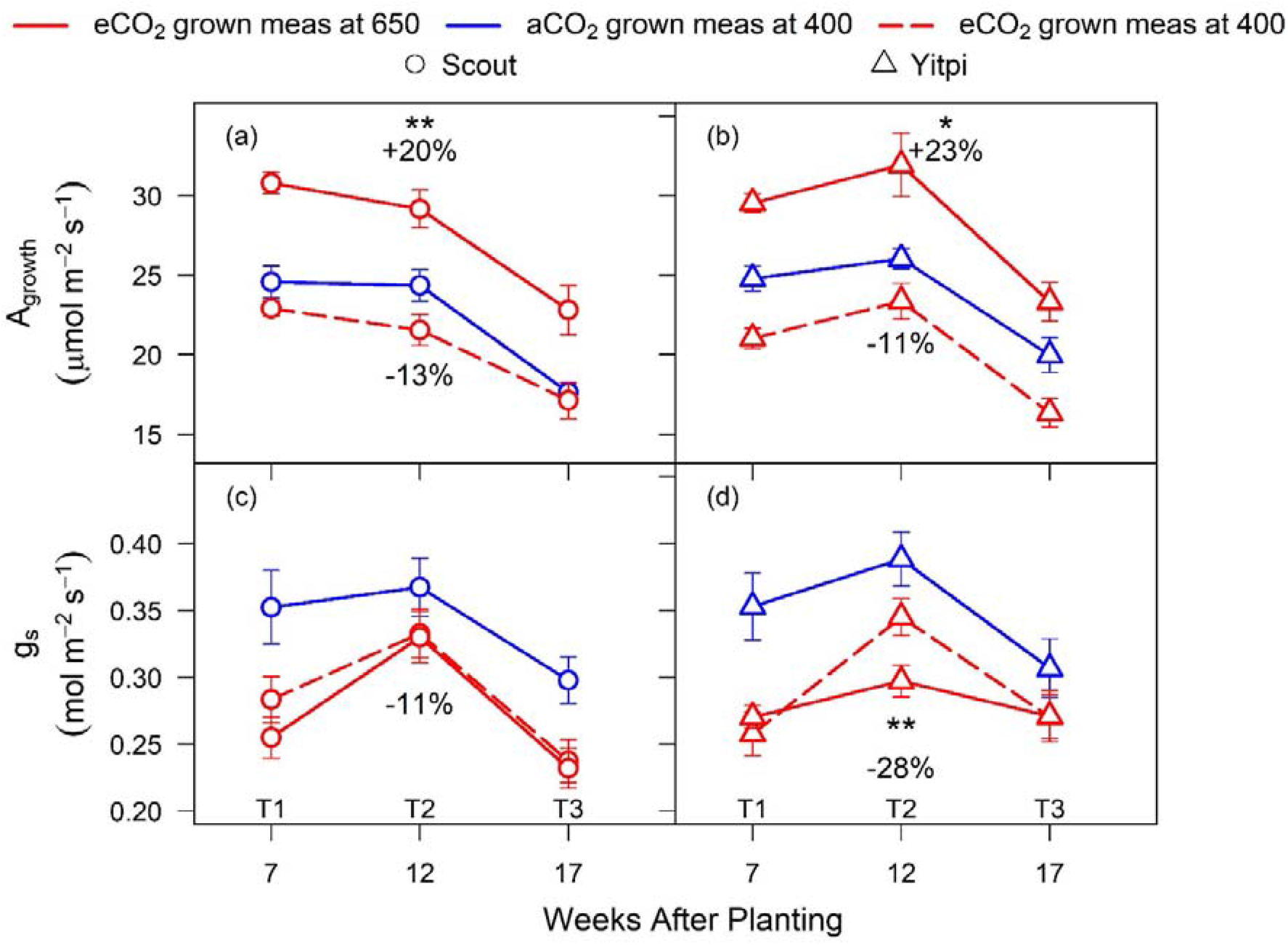
Response of leaf gas exchange parameters to eCO_2_ under non-HS conditions. Measurements were made at 25°C before each harvest (T1, T2 and T3) for CO_2_ assimilation rates (a, b) and stomatal conductance (c, d) in Scout (Circles) and Yitpi (Triangles). Plants were grown and measured at aCO_2_ (blue solid cultivars), grown and measured at eCO_2_ (red solid cultivars), and grown at eCO_2_ and measured at 400 μL CO_2_ L^-1^ (red dashed cultivars). Statistical significance levels (t-test) for the growth condition within each cultivar are shown and they are: * = *p* < 0.05; ** = *p* < 0.01: *** = *p* < 0.001.

### High temperature during HS enhanced photosynthesis under eCO_2_

The two HS cycles did not reduce *A_growth_* measured at 25℃ during or after HS (Figure 3A-D, Tables S1-S3). During both HS1 and HS2, eCO_2_ stimulated *A_growth_* measured at 25℃ in Scout but not Yitpi. Relative to 25°C, *A_growth_* increased at 35°C in Scout (10-14%) and Yitpi (12-18%) grown at eCO_2_ but not at aCO_2_. Immediately after the recovery from HS, *A_growth_* was upregulated in eCO_2_-grown Scout (Figures 3A-D and S3). During both HS cycles, dark-adapted Fv/Fm measured at 25℃ tended to be lower in Yitpi grown at eCO_2_ relative to aCO_2_. In both cultivars, Fv/Fm decreased at 35℃ relative to 25°C, indicating transient damage to PSII due to HS at both CO_2_ treatments (Figure 3E-H, Tables S1-S3).

**Figure 3.**
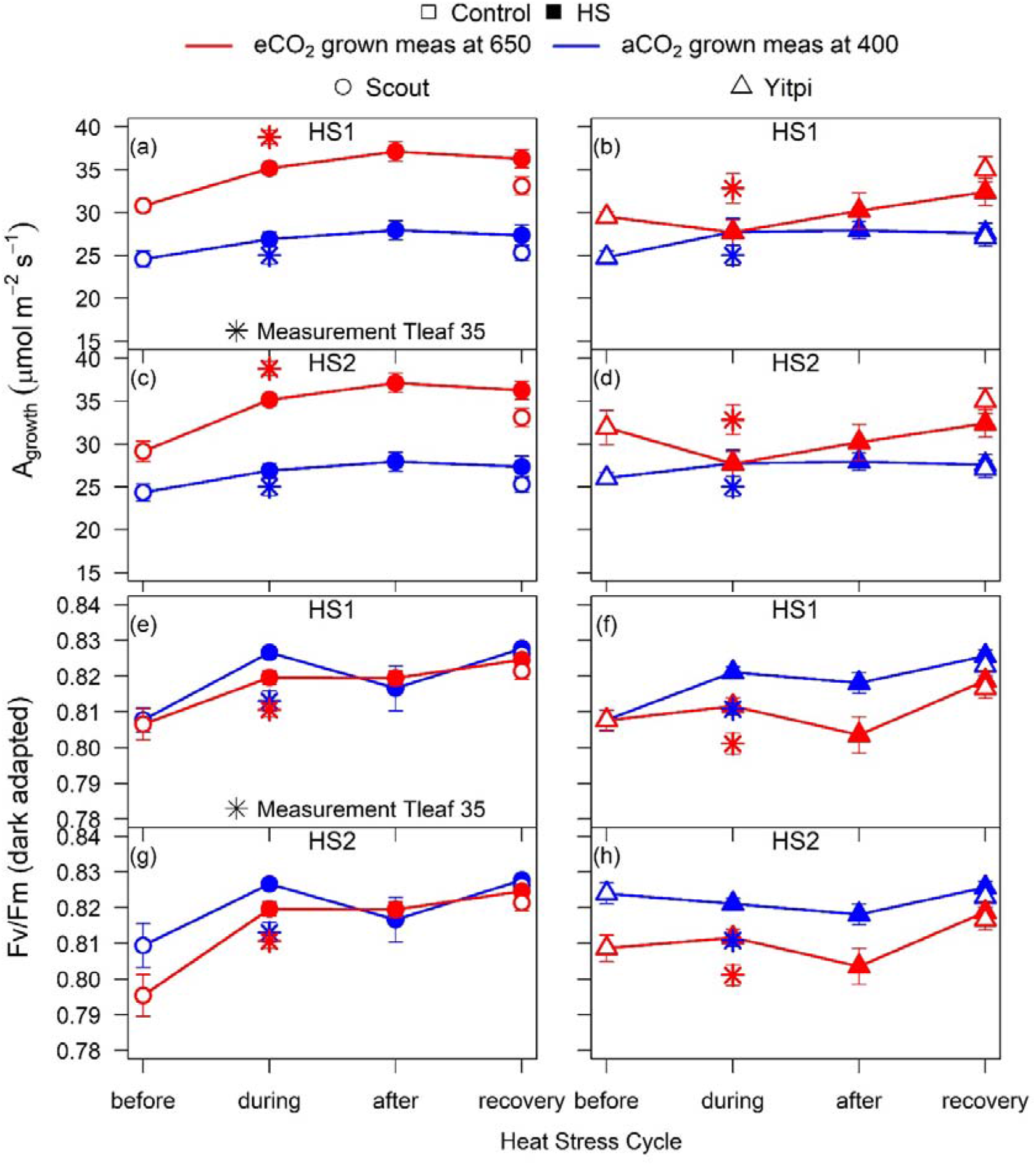
Response of photosynthesis and chlorophyll fluorescence to HS in Scout and Yitpi grown at aCO_2_ or eCO_2_. CO_2_ assimilation rates (a, b, c, d) and dark-adapted chlorophyll fluorescence, Fv/Fm (e, f, g, h) were measured at growth CO_2_ and 25 °C in Scout (Circles) and Yitpi (Triangles). Open and closed symbols represent control and HS plants, respectively. In addition, plants were measured at 35°C (*) during both HS cycles.

Following long-term recovery from HS1 and/or HS2, the eCO_2_ stimulation of *A_growth_* was still marginally apparent in all T3 plants, being the strongest in eCO_2_-grown Yitpi (Figure 4A-B, Tables S1-S3). The reduction of *g_s_* at eCO_2_ was weak in all plants (Figure 4C-D, Tables S1-S3). Hence, PWUE was stimulated by eCO_2_ in all treatments, while PNUE was unaffected (Figure 4E-H, Tables S1-S3). There was a good correlation between *A_growth_* and *g_s_* (*r*^2^ = 0.51, p <0.001) across all treatments (Figure 5A).

**Figure 4.**
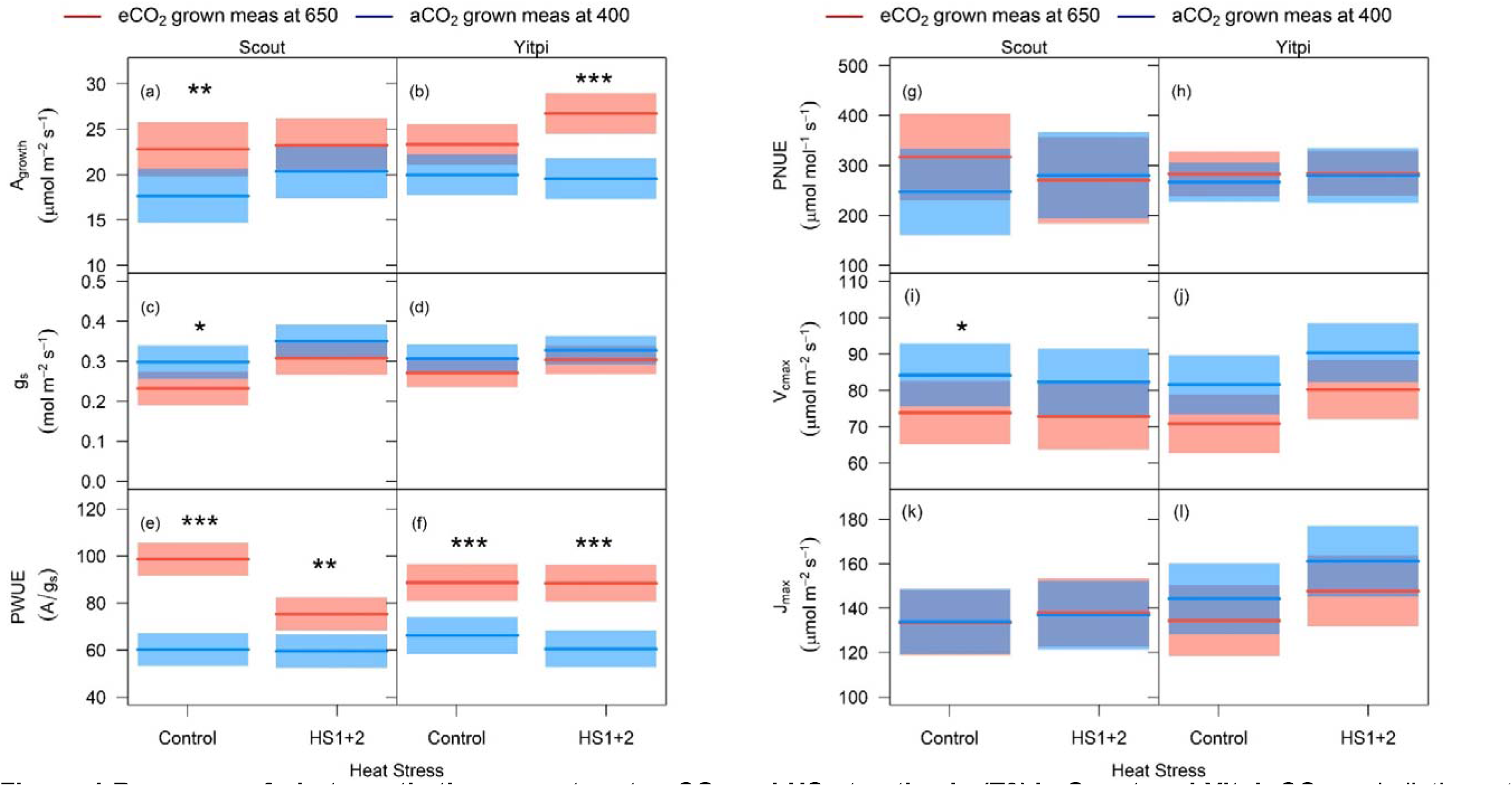
Response of photosynthetic parameters to eCO_2_ and HS at anthesis (T3) in Scout and Yitpi. CO_2_ assimilation rate (a, b), stomatal conductance (c, d), photosynthetic water use efficiency (e, f) and photosynthetic nitrogen use efficiency (g, h) were measured at growth CO_2_. *V_cmax_* (i, j) and *J_max_* (k, l) were derived from ACi curves measured at 25°C. Cultivars indicate means and shaded region is 95% confidence interval. Data shown for control (not exposed to any heat stress) and plants exposed to both heat stress cycles (HS1+2). Statistical significance levels (t- test) for the growth condition within each cultivar are shown and they are: * = *p* < 0.05; ** = *p* < 0.01: *** = *p* < 0.001.

**Figure 5.**
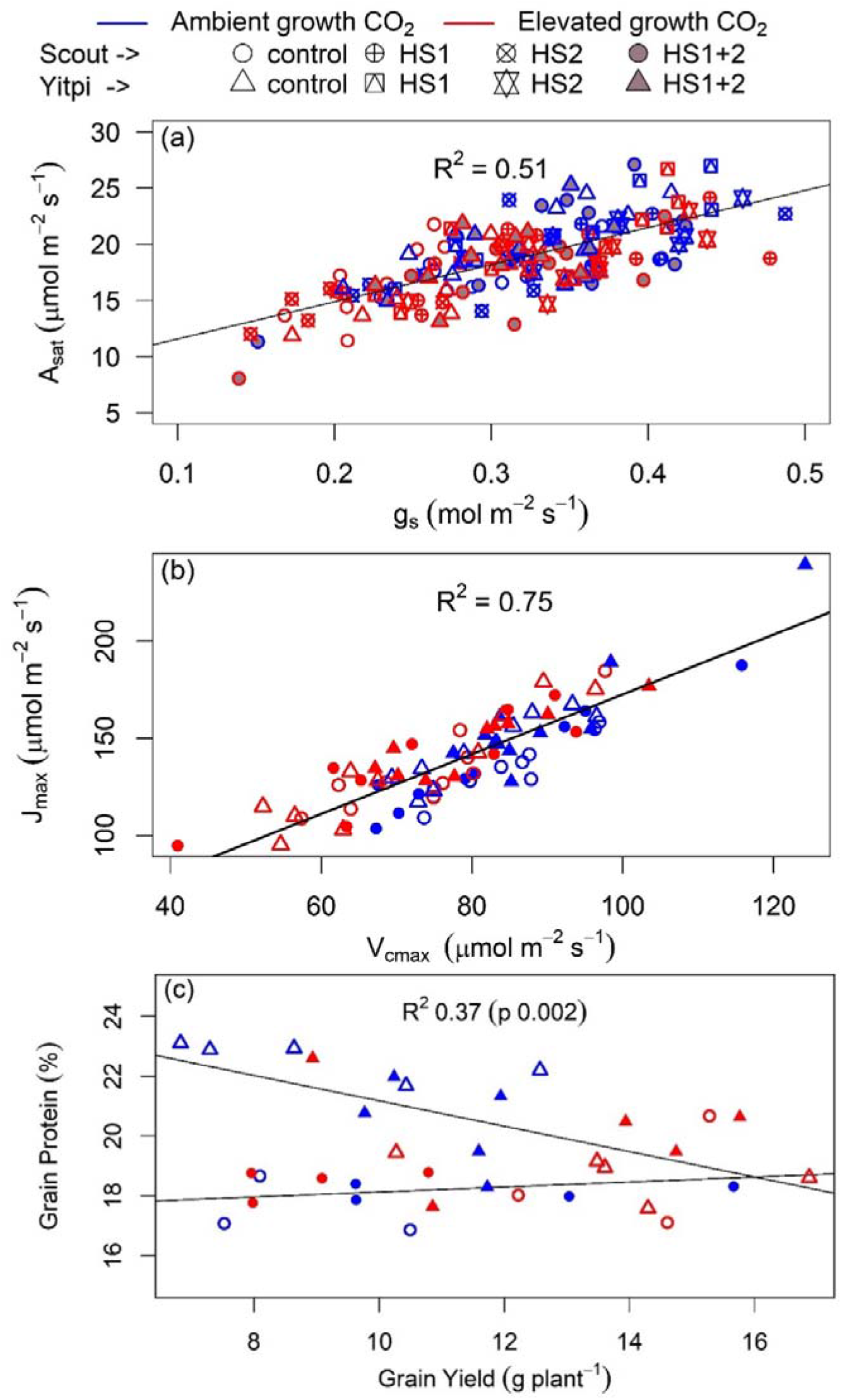
Relationships with leaf gas exchange and grain yield across treatments. CO_2_ assimilation rate plotted as a function of stomatal conductance (a) (both aCO_2_ and eCO_2_ grown plants measured at 400 μl L^-1^), *J_max_* plotted as a function of *V_cmax_* (b) and grain protein plotted as a function of yield (c) in Scout (Circles) and Yitpi (Triangles). Ambient and elevated CO_2_ are depicted in blue and red, respectively. Control and heat stressed plants depicted using open and closed symbols. Panel a depicts data for control, HS1, HS2 and both heat stresses (HS1+2), while panels b and cinclude only control and HS1+2.

*V_cmax_* and *J_max_* were derived from A-C_i_ response curves measured at 25°C during the recovery stage after HS2. For control and HS plants, growth at eCO_2_ marginally reduced *V_cmax_* in Scout (−14%, p = 0.09) and Yitpi (−15%, p = 0.06) but had no effect on *J_max_*. HS had no effect on *V_cmax_* or *J_max_* in either cultivar (Figure 4I-L, Tables S1-S3). *V_cmax_* and *J_max_* correlated well (*r*^2^ = 0.75, p <0.001) across treatments (Figure 5B).

### Yitpi produced more tillers and grains than Scout

When compared at aCO_2_, the two wheat cultivars differed in phenology and growth habit. Scout developed faster and flowered earlier than Yitpi. At T2, 43% of tillers had ears in Scout compared to 11% in Yitpi (Figure S4). At T2, Scout was 74% (p < 0.001) taller than Yitpi but at T3 both cultivars had similar height (Figure 6I-J, Tables S4 and S5). In contrast, Yitpi accumulated more biomass relative to Scout by producing more tillers. At T3, Yitpi had 42% (p < 0.005) more total plant biomass, 130% (p < 0.001) more tillers, 254% (p < 0.001) larger leaf area, 128% (p < 0.001) more leaves and 61% (p < 0.001) larger leaf size compared to Scout (Figure 6, Tables S4 and S5).

**Figure 6.**
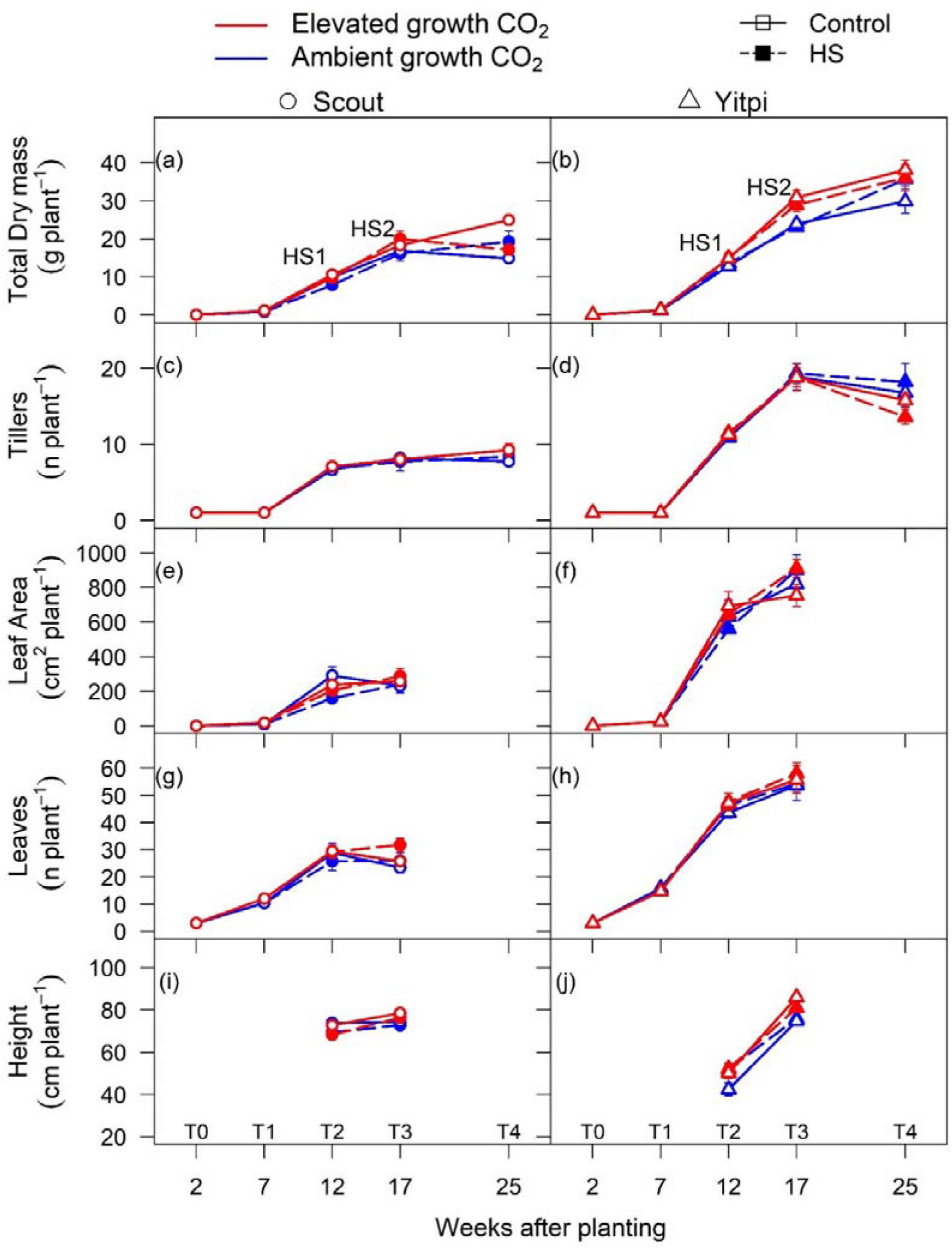
Response of plant growth and morphological traits to elevated CO_2_ and HS: Total dry mass (a, b), tillers or number of tillers (c, d), leaf area (e, f), leaf number (g, h) and height (i, j) were measured at different time points across the life cycle of wheat cultivars Scout (Circles) and Yitpi (Triangles). Ambient and elevated CO_2_ are depicted in blue and red color, respectively. Open symbols connected with solid cultivars and closed symbols connected with dashed cultivars represent control and HS plants, respectively. HS1 and HS2 depict the timing of HS applied at 10 and 15 weeks after planting respectively.

At the final harvest (T4), Yitpi had more plant biomass (84%, p < 0.001), tillers (88%, p < 0.001) and number of grains (54%, p < 0.001). Conversely, Scout had larger grain size (+31%, p < 0.001), a higher proportion (100%) of its tillers developed ears and more ears filled grains compared to higher tillering Yitpi (88%). Hence, both cultivars had relatively similar grain yield (g/plant) (Figure 7A-F, Tables S4 and S6). Higher (178%, p < 0.001) harvest index (HI) in Scout was due to early maturity and consequent leaf senescence leading to loss of biomass at final harvest (Tables S5). The final harvest was undertaken four weeks after all ears had matured on Scout to give ample time for grain filling in Yitpi (Figure 7, Tables S4 and S6).

**Figure 7.**
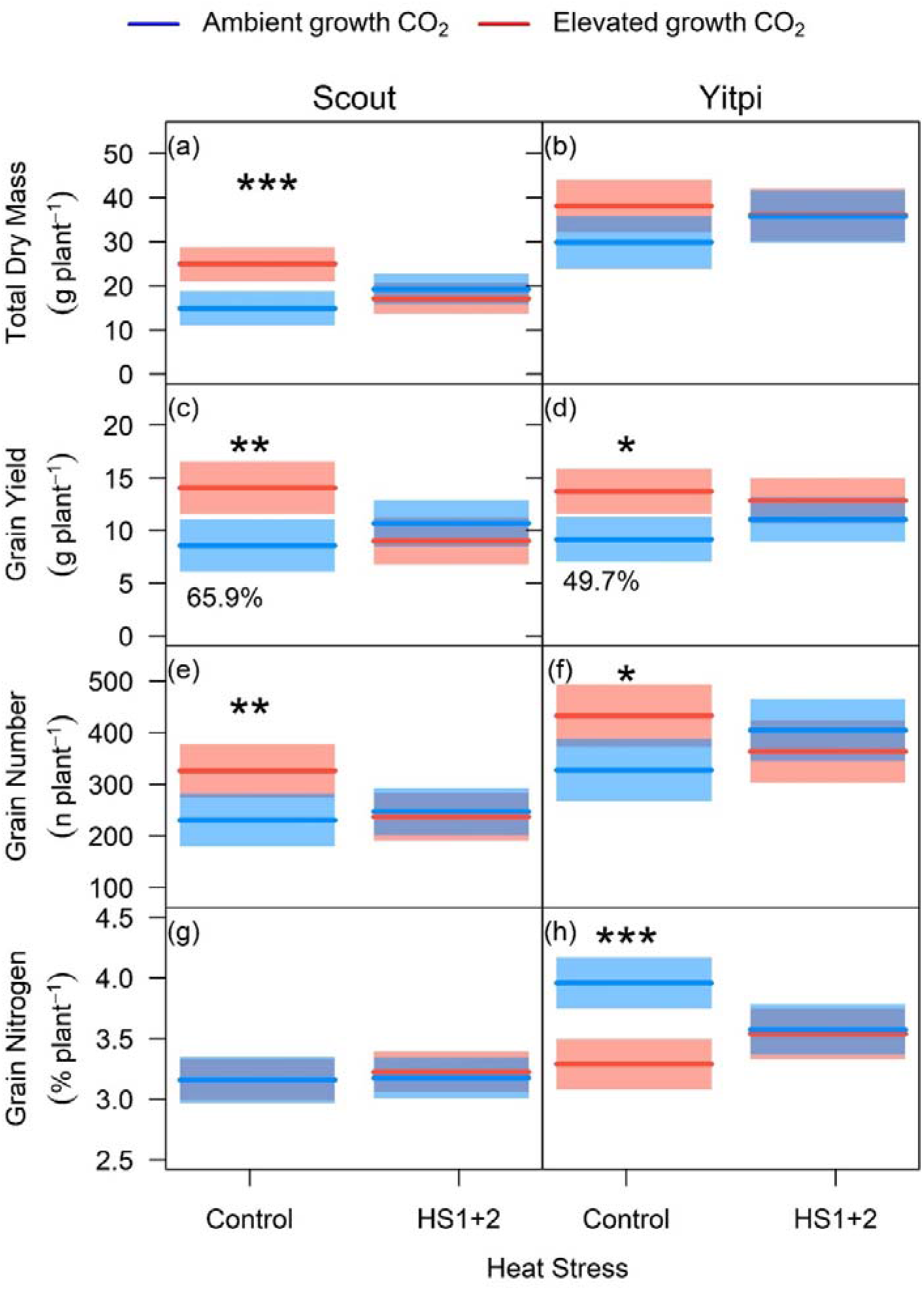
Response of total plant dry mass and grain parameters to growth at eCO_2_ and HS at maturity (T4): Total dry mass (a, b), grain dry mass (c, d), grain number (e, f) and grain nitrogen (g, h) were measured at the final harvest. Cultivars indicate means and shaded region is 95% confidence interval. Ambient and elevated CO_2_ are depicted in blue and red color respectively. Heat stress levels include plants not exposed to any heat stress (control) and both heat stresses (HS1+2). Statistical significance levels (t-test) for the growth condition within each cultivar is shown and they are: * = *p* < 0.05; ** = *p* < 0.01: *** = *p* < 0.001.

### eCO_2_ similarly stimulated wheat biomass and grain yield under non-HS conditions

The increase in plant biomass at eCO_2_ depended on the growth stage (Figures 6-7, Tables S4 and S5). However, the overall stimulation was not different between the two cultivars as evident from the non-significant eCO_2_ x cultivar interaction at all harvests (Table S4). By T3 (anthesis), when both cultivars were still within the exponential growth stage, eCO_2_ stimulated plant biomass of Yitpi (+29%, p < 0.001) and Scout (+9%, p < 0.001) under control conditions. The number of tillers, total leaf area, mean leaf size or leaf mass area were not significantly affected by eCO_2_ in either cultivar (Figure 6, Tables S4 and S5). eCO_2_ increased allocation to stem relative to leaf biomass, particularly in Yitpi. Accordingly, there was a strong correlation across treatments between stem and leaf biomass (*r^2^ = 0.83*, p < 0.001) and between total biomass and leaf area (*r^2^ = 0.83*, p < 0.001) in Scout but not in Yitpi. However, the two cultivars followed common relationship for root *versus* shoot biomass (*r^2^ = 0.41,* p < 0.001) and leaf area *versus* leaf number (*r^2^ = 0.82*, p < 0.001) across all treatments suggesting no effect of cultivar, eCO_2_ or HS on these common allometric relationships (Figure S5, Table S5).

At the final harvest T4 (seed maturity), eCO_2_ enhanced biomass and equally stimulated grain yield by increasing grain number in both cultivars (+64% in Scout and +50% in Yitpi) under control conditions only (Figure 7A-D, Tables S4-S6). Harvest index was not directly affected by any treatments but showed a significant interaction (p <0.05) between CO_2_ and cultivar, such that HI was higher in Yitpi under eCO_2_ (Tables S5 and S6).

### eCO_2_ did not stimulate the grain yield of HS plants

At T3, moderate HS (34-38°C) applied under well-watered conditions and 60% RH during the vegetative (HS1 applied after T1) and flowering (HS2 applied after T2) stages had no significant impact on plant biomass of either wheat cultivar or CO_2_ treatment. By T4, there were significant HS x CO_2_ x cultivar interactions (p < 0.01) for biomass and grain yield. HS1+2 reduced the biomass and grain yield of eCO_2_- grown Scout relative to aCO_2_-grown counterparts. Unlike control plants, the biomass and yield of HS plants were not enhanced by eCO_2_ (Figure 7, Tables S4 and S5).

### eCO_2_ reduced grain N in Yitpi but not in Scout

Neither eCO_2_ or HS had a significant effect on flag leaf N content in either cultivar at T2 or T3, but eCO_2_ reduced aggregate leaf N content (−18%) at T3 in Yitpi only (Cultivar x CO_2_ p < 0.05) (Table S7). Yitpi had higher grain N content (+26%) than Scout in control plants grown at aCO_2_ (Figure 7G-H, Table S6). In control plants, eCO_2_ significantly reduced grain protein content in Yitpi (−18%, p < 0.05) but not in Scout due to significant cultivar X CO_2_ interaction (p < 0.01), while HS had no effect on protein content in either cultivar (Figure 7G-H, Table S6).

## Discussion

### Two wheat cultivars with contrasting morphology and phenology, but similar photosynthesis and grain yield

The effects of future climate conditions, including eCO_2_, will depend on the environmental conditions (e.g., water and heat stress) and the crop’s agronomic features. Here, we compared the interactive effects of eCO_2_ and HS on two commercial wheat cultivars, Scout and Yitpi, with contrasting phenology and growth habit. Plants were grown under well-watered and fertilized conditions to remove any confounding effects of water or nutrient limitations on the eCO_2_ or HS responses. RH was kept constant to minimize the negative impact of dry air during HS. Finally, we compared the effects of applying HS at the vegetative and flowering stages.

Free tillering Yitpi produced substantially more tillers, leaf area and biomass relative to the faster developing Scout. Accordingly, our first hypothesis predicted that Yitpi will have higher grain yield. The results only partially supported this hypothesis, because relative to Yitpi, Scout had higher harvest index (HI) due to its early maturing and senescing habit. While Yitpi initiated more tillers, a lower proportion of these tillers produced ears and filled grains. In contrast, Scout produced less tillers but flowered earlier which allowed enough time for all its tillers to produce ears and fill bigger grains by the final harvest. Hence, both cultivars had relatively similar yields due to bigger grain size in Scout and higher grain number in Yitpi. It is worth noting that some field trials have reported slightly higher grain yields in Scout than Yitpi (National variety trial report, GRDC, 2014). Our results are consistent with a previous study using different wheat cultivars with contrasting source-sink relationships which reported that the freely tillering cultivar “Silverstar” translated into more spikes while restricted tillering cultivar “H45” had more and heavier kernels per spike than “Silverstar” (Tausz-Posch et al., 2015). Thus, early vigor and maturity compared to high tillering capacity seem to be equally beneficial traits for high grain yield in the Australian environment.

The two wheat cultivars showed similar photosynthetic traits and response to temperature and eCO_2_. In contrast to our expectations that Scout would have higher WUE due to its selection based on a carbon isotope discrimination gene (Condon *et al*., 2004), both wheat cultivars showed similar PWUE under most measurement and growth conditions in this study (Figure 4E-F, Table 2).

### Elevated CO_2_ stimulated photosynthesis but reduced photosynthetic capacity in both cultivars

Long term exposure to eCO_2_ may reduce photosynthetic capacity due to lower amount of Rubisco in a process referred as ‘acclimation’ (Nie *et al*., 1995; Rogers and Humphries, 2000; Ainsworth *et al*., 2003). Alternatively, eCO_2_ may ‘down- regulate’ photosynthetic capacity by reducing Rubisco activation or other regulatory mechanisms without affecting Rubisco content (Delgado *et al*., 1994). In the current study, eCO_2_ similarly increased *A_growth_* (+21%) measured at growth CO_2_ and reduced both *A_sat_* (−12%) measured at common CO_2_ (Figure 2, T2) and *V_cmax_* (Figure 4I-J) in both cultivars. In contrast, leaf N (and possibly Rubisco) was not significantly affected in either cultivar (Table S4). Hence, the wheat cultivars have likely undergone a photosynthetic downregulation -rather than acclimation- in response to eCO_2_ (Delgado *et al*., 1994; Leakey *et al*., 2009). These results partially countered our second hypothesis suggesting that Yitpi will show less photosynthetic acclimation due to its higher sink capacity. The interaction of eCO_2_ with plant traits are complex. On the one hand, eCO_2_ is expected to cause less feedback inhibition on photosynthesis in plants with high sink capacity (Ainsworth *et al*., 2004). On the other, fast-growing plants show a proportionally larger response to eCO_2_ (Poorter and Navas, 2003). Hence, high tillering in Yitpi and fast development in Scout both led to a relatively small observed photosynthetic downregulation in response to growth at eCO_2_. This allowed a sustained photosynthetic stimulation, which in turn led to a significant biomass and yield enhancement by CO_2_ enrichment in both wheat cultivars (Figures 6-7). Photosynthetic responses of wheat in current study are in agreement with earlier enclosure studies which generally have higher response to eCO_2_ than the FACE studies (Kimball *et al*., 1995; Hunsaker *et al*., 1996; Osborne *et al*., 1998; Kimball *et al*., 1999; Long *et al*., 2006; Cai *et al*., 2016).

### Elevated CO_2_ stimulated grain yield similarly in both wheat cultivars

In disagreement with our second hypothesis, eCO_2_ similarly stimulated plant biomass and grain yield in early-maturing Scout and high tillering Yitpi (Figures 6-7, Table S4-S5). In Scout, the biomass stimulation was associated with increased tillering (one extra tiller per plant). In contrast, Yitpi produced many tillers at aCO_2_ and the additional fixed carbon at eCO_2_ was allocated to the existing tillers. At seed maturity, eCO_2_ stimulated grain yield similarly in both cultivars as a result of the trade-off between grain yield components (Dias de Oliveira *et al*., 2015). In particular, eCO_2_ stimulated grain number in both cultivars, while grain size increased in Scout only (Figure 7, Table S6). Generally, eCO_2_ stimulates grain yield by increasing the number of tillers and consequently, ears per plant (Zhang *et al*., 2010; Bennett *et al*., 2012), which has also been reported in FACE studies (Högy *et al*., 2009; Tausz-Posch *et al*., 2015; Fitzgerald *et al*., 2016). However, in our study, the increase in grain yield at eCO_2_ was mainly due to the increase in the number of grains per ear. In line with our results, Tausz-Posch et al., (2015) reported comparable grain yield stimulation by eCO_2_ in two different wheat cultivars with contrasting source-sink relationships. Moreover, grain yield of twenty wheat cultivars that differed in tillering propensity, water soluble carbohydrate accumulation, early vigor and transpiration efficiency responded similarly to eCO_2_ in glasshouse settings (Ziska *et al*., 2004; Bourgault *et al*., 2013).

### Elevated CO_2_ reduced grain N in Yitpi only

Overall, there is a negative relationship between grain yield and quality (Taub *et al*., 2008; Pleijel and Uddling, 2012). Hence, increased grain yield at eCO_2_ results in lower grain N and hence protein content (Seneweera and Conroy, 1997; Bahrami *et al*., 2017). In our study, eCO_2_ reduced grain N in Yitpi under control conditions. Scout was characterized by having larger grains which accumulated less N than Yitpi. Moreover, eCO_2_ reduced total leaf N (−18%) at T3 and grain N (−17 %) at T4 in Yitpi but not in Scout. This is consistent with the results from FACE study with same cultivars which reported -14% reduction in N content by eCO_2_ in above ground dry mass in Yitpi but not in Scout under well-watered conditions (Bahrami *et al*., 2017). The higher biomass accumulation in free tillering Yitpi may have exhausted the nutrient supply, such that further biomass stimulation by eCO_2_ lead to a significant dilution in N content (Taub and Wang, 2008).

Wheat cultivars with early vigour such as Scout have greater root biomass accumulation as well as greater early N uptake which may have avoided a negative effect of eCO_2_ on leaf and grain N (Liao *et al*., 2004; Bahrami *et al*., 2017). Accordingly, Scout maintained a higher N utilization efficiency (grain yield per total plant N) relative to Yitpi under all treatments (Table S6). Increased grain yield is strongly associated with higher grain number per unit area (Zhang *et al*., 2010; Bennett *et al*., 2012) which dilutes the amount of N translocated per grain. Quality deterioration due to lower protein via reduced N is of critical concern in future high CO_2_ climate considering that even additional supply of N does not prevent N dilution in grain under eCO_2_ (Tausz *et al*., 2017). In addition, eCO_2_ has strong detrimental effect on other nutrient availability and remobilization from leaves to grains (Tcherkez *et al*., 2020).

### HS had little effects on wheat photosynthesis or yield at aCO_2_

A key finding of this study was that the application of HS events (HS1, HS2 or HS1+2) was not detrimental to aCO_2_-grown wheat plants (Figures 5 and 7, Tables S4-S6). Thus, our hypothesis that HS will reduce photosynthesis, biomass and yield at aCO_2_ was rejected. This finding is in contrast to previously reported studies where HS reduced the grain yield and negatively affected the growth and development in wheat (Stone and Nicolas, 1996, 1998; Farooq et al., 2011; Coleman et al., 1991). During heat waves in the field, the vapor pressure deficit (VPD) increases and soil moisture decreases leading to lower stomatal conductance and consequently lower transpiration rate. Thus, plants are unable to cool down and leaf temperatures rise beyond optimum levels causing damage. The negligible effect of HS in our study could be explained by the ability of well-watered plants to maintain leaf temperature below damaging levels due to transpirational cooling (Perera *et al*., 2019; Deva *et al*., 2020) even with air temperatures reaching up to 38°C. At moderate (∼60%) relative humidity, there is sufficient water vapour gradient to sustain high transpiration rates when soil water is available, as was the case in our experiment. In most cases, *g_s_* was not significantly affected (Tables S1 and S2), and even slightly higher at T3 in HS-pants relative to the control (Figure 4c,d) Well-watered crops can maintain grain-filling rate, duration and size under HS (Dupont *et al*., 2006), and high temperatures can increase crop yields if not exceeding critical optimum growth temperature (Welch *et al*., 2010). Also, in the current study, the night temperatures were not increased during HS which favors plant growth by reducing respiratory losses (Prasad *et al*., 2008).

In particular, HS did not elicit a direct negative impact on photosynthesis or chlorophyll fluorescence in either cultivar or CO_2_ treatment. During HS, high temperature transiently reduced maximum efficiency of PSII (Fv/Fm) in both cultivars and CO_2_ treatments (Figure 3E-H). However, unchanged Fv/Fm measured at 25°C confirmed that photosynthesis did not suffer long-term damage during or after HS. Moreover, HS was not severe enough to negatively affect *A_growth_* measured at 25°C. These results are corroborated by the insensitivity of *V_cmax_* and *J_max_* to HS (Figure 4I- L), but contrast with previously reported studies where HS reduced photosynthesis in wheat at the vegetative (Wang et al., 2008) and the flowering (Chavan *et al*., 2019; Balla *et al*., 2019) stages. HS lowers membrane thermostability by inducing reactive oxygen species (ROS) and altering the membrane protein structures, which lead to changes in the fluidity of the thylakoid membrane and separation of light harvesting complex from the photosystems (Wahid *et al*., 2007; Poudel, 2020). We were unable to measure leaf temperatures in the current study, but we speculate that, in well-watered wheat plants growing at moderate RH, leaf temperatures might not have increased beyond damaging levels to the membranes during the HS events.

Repeated HS may result in priming which involves pre-exposure of plants to a stimulating factor such as HS (Wang *et al*., 2017) and enable plants to cope better with later HS events (Balla *et al*., 2021). However, there was no difference between HS applied at the vegetative (HS1) and/or flowering stage (HS2) in rejection of our fourth hypothesis, and this may additionally be due to the short term duration of the two HS cycles (3 days each). Hence, our study demonstrated the benign effect that HS has on crop yield when separated from water stress and plants are able to transpire.

### HS precluded an eCO_2_ response in biomass and grain yield

In our study, the impact of HS depended on the wheat cultivar and growth CO_2_ (Tables S1 and S4). Elevated CO_2_ and temperature interactions can be complex, dynamic and difficult to generalize as they can go in any direction depending on plant traits and other environmental conditions (Rawson, 1992). Plant development is generally accelerated by increased temperature; eCO_2_ can accelerate it further in some instances or may have neutral or even retarding effects in other cases (Rawson, 1992).

While eCO_2_ stimulated wheat biomass and grain yield under control (non-HS) conditions, HS precluded a yield response to eCO_2_ in Yitpi and reduced biomass and yield in eCO_2_-grown Scout relative to aCO_2_-grown counterparts (Figure 7). These results are in contrast with previous studies that reported similar wheat yield reduction at ambient or elevated CO_2_ in response to severe (Chavan *et al*., 2019) or moderate HS (Zhang *et al*., 2018). The results also partially refuted our third hypothesis that HS may decrease yield more at aCO_2_ than eCO_2_, while partially agreeing that HS will have a more negative impact on Scout relative to Yitpi, albeit for different reasons than what we originally suggested. The negative effect of HS on Scout biomass and grain yield at eCO_2_ occurred despite the eCO_2_ stimulation of *A_growth_* under HS (T3, Figure 4). However, over the long term, *A_growth_* was stimulated in eCO_2_-grown Yitpi and not Scout (Figure 4).

Lack of a biomass stimulation despite high photosynthetic rates during HS under eCO_2_ could be due to the short duration of HS (3 days), which may not have been long enough to stimulate biomass gain. In addition, nutrient limitation at eCO_2_ may have restricted the eCO_2_ growth response. Typically, eCO_2_ studies show reduced N content in wheat and other crops (Taub and Wang, 2008; Leakey *et al*., 2009; Bahrami *et al*., 2017). Hence, the wheat plants may have exhausted available nutrients due to increased demand by growing sinks at eCO_2_, which may limited further stimulation by high temperature. HS may be more damaging at eCO_2_ due to reduce transpirational cooling as a result of reduced g_s_ at eCO_2_, leading to higher leaf temperatures. However, *A_growth_* increased in response to high temperature (35°C) under eCO_2_ but not under aCO_2_ during HS (Figure 3A-D), which refutes the suggestion of HS-damage to photosynthesis.

Higher g_s_ during HS at moderate RH in well-watered conditions may increase *A_growth_* by increasing C_i_ in both aCO_2_ and eCO_2_ grown plants. Furthermore, lower photorespiration under eCO_2_ allows additional increase in *A_growth_* with temperature when measured at 35°C relative to 25°C (Long, 1991). Under aCO_2_, photorespiration increases with temperature reducing *A_growth_* measured at 35°C relative to 25°C. Our results also point to a shift in *T_opt_* of photosynthesis (∼ 24°C at aCO_2_) to higher temperatures for plants grown at eCO_2_ (Sage and Kubien, 2007). This would come about as a result of lower photorespiration at eCO_2_ as well as the slight upregulation of photosynthetic rates observed in eCO_2_-grown Scout at the recovery stage of HS (Figure 3A, C). However, at T3, *A_growth_* was similar between aCO_2_ and eCO_2_ grown plants (Figure 4A) indicating the short-term nature of this photosynthetic upregulation.

## Conclusions

The two wheat cultivars, Scout and Yitpi differed in growth and development but produced similar grain yield. Under control conditions, eCO_2_ stimulated biomass and yield similarly in both cultivars. HS was not damaging to photosynthesis, growth, biomass or grain yield under well-watered and moderate RH conditions. However, HS interacted with eCO_2_, leading to similar or lower biomass and grain yield at eCO_2_ relative to both aCO_2_ in plants exposed to HS. This interactive effect precluded the positive effects of eCO_2_ in HS-plants. eCO_2_ improved photosynthetic rates in control and HS plants. Also, high temperature stimulated photosynthesis under eCO_2_ but not under aCO_2_ during HS which suggests increased optimum temperature of photosynthesis at eCO_2_. We speculate that in the current study, HS plants were able to cool down using high transpiration which helped to maintain lower leaf temperatures despite high air temperatures during HS. The current study provides important insights into the effect of short-term moderate temperature increases in well-watered conditions under future elevated CO_2_, potential role of transpirational cooling during HS and interactions between HS and eCO_2_ which will be useful in breeding cultivars for future climate and improving crop model accuracy to predict crop performance in future high CO_2_ environment with frequent heat waves.

## Acknowledgments

We gratefully acknowledge the technical support of Fiona Koller. SGC was supported by the ‘Agriculture, Fisheries & Forestry Postgraduate Research Scholarship’ and Western Sydney University. The research received funding support by the Australian Commonwealth Department for Agriculture and Water Resources through the ‘Filling the research gap’ programme and was associated with the Australian Grains Free Air CO_2_ Enrichment (AGFACE) programme run jointly by The University of Melbourne and Agriculture Victoria. OG was also funded by the Australian Research Council Centre of Excellence for Translational Photosynthesis (CE140100015).

## Availability of data

All data supporting the findings of this study are available within the paper and within its supplementary materials published online. Reuse of the data is permitted after obtaining permission from the corresponding author.

## Code availability

Software application and codes used are all publicly available.

## Authors contributions

All authors conceived the project. SGC maintained the plants and collected the data. SGC and RAD analysed the data. SGC and OG prepared the manuscript with input from other co-authors.

## Supplementary Materials

**Table S1.** Summary of statistics for gas exchange parameters.

**Table S2.** Response of Scout gas exchange parameters to elevated CO_2_ and heat stress.

**Table S3.** Response of Yitpi gas exchange parameters to growth at elevated CO_2_ and heat stress.

**Table S4.** Summary of statistics for plant dry mass and morphological parameters.

**Table S5.** Response of plant dry mass and morphological parameters to elevated CO_2_ and HS.

**Table S6.** Summary of plant nitrogen content parameters.

**Figure S1.** Glasshouse growth conditions.

**Figure S2.** Experimental design.

**Figure S3.** Photosynthetic response to growth at eCO2 and heat stresses (HS1 and HS2).

**Figure S4.** Response of ear development in Scout and Yitpi to eCO_2_ and HS at the booting stage (T2).

**Figure S5.** Relationship between dry mass and morphological parameters measured at anthesis (T3).

